# MultiVI: deep generative model for the integration of multi-modal data

**DOI:** 10.1101/2021.08.20.457057

**Authors:** Tal Ashuach, Mariano I. Gabitto, Michael I. Jordan, Nir Yosef

**Affiliations:** Center for Computational Biology, University of California, Berkeley, CA, USA; Department of Electrical Engineering and Computer Sciences, University of California, Berkeley, CA, USA; Department of Statistics, University of California, Berkeley, Berkeley, CA, USA; Ragon Institute of MGH, MIT, and Harvard, Cambridge, MA, USA; Chan Zuckerberg BioHub, San Francisco, CA, USA; Allen Institute for Brain Science, Seattle, WA, USA

**Author notes:** Contributed Equally (Tal Ashuach, Mariano I. Gabitto).

## Abstract

Jointly profiling the transcriptional and chromatin accessibility landscapes of single-cells is a powerful technique to characterize cellular populations. Here we present MultiVI, a probabilistic model to analyze such multiomic data and integrate it with single modality datasets. MultiVI creates a joint representation that accurately reflects both chromatin and transcriptional properties of the cells even when one modality is missing. It also imputes missing data, corrects for batch effects and is available in the scvi-tools framework: https://docs.scvi-tools.org/.

## 1 Introduction

The advent of technologies for profiling the transcriptional and chromatin accessibility landscapes at a single cell resolution has been paramount for cataloging cellular types and states, identifying important genomic regions, and linking genes to their regulatory elements [1, 2]. However most uses of single-cell RNA-seq (scRNA-seq) [3, 4] and single-cell ATAC-seq (scATAC-seq) [2, 5] have been limited such that a given cell can only be profiled by one technology. Recently, multi-modal single-cell protocols [6] have emerged for simultaneously profiling gene expression and chromatin accessibility in the same cell. This concomitant measurement is promising to enable a more refined categorization of cell states and, ultimately, a better understanding of the regulatory mechanisms that underlie the diversity of states.

The emerging area of multi-modal profiling has benefited greatly from new statistical methods that jointly account for both modalities in a range of analysis tasks [7, 8]. Another promising application of multi-modal assays, however, is to improve the way by which the more common and less costly single-modality datasets (profiled with scRNA- or scATAC-seq) are analyzed and interpreted. By leveraging the paired information, one can infer properties of the missing modality and thus derive new insight about the diversity of cell states and the regulation of gene expression in these datasets. To provide a comprehensive solution, such an integrative analysis should be done at two levels. First, it will generate a low dimensional summary of the state of each cell that reflects both its transcriptional and chromatin properties, regardless of whether the cell was profiled with one or both modalities. As commonly done in other applications of single cell genomics, such a representation can facilitate the identification of sub-populations or gradients and enable data visualization [**vision**]. A second level of analysis will generate a normalized, batch corrected view of the high-dimensional data (gene expression, chromatin accessibility), either observed or inferred. Such an analysis can enable the identification of loci or genes whose accessibility or expression characterize sub-populations of interest.

Here, we introduce MultiVI, a deep generative model for probabilistic and integrative analysis of scRNA-seq, scATAC-seq and paired multi-modal data. MultiVI operates at the two levels of analysis, providing both a low-dimensional summary of cell state and a normalized high dimensional view of gene expression and chromatin accessibility. MultiVI was designed to account for the general caveats of single cell genomics data, namely: batch effects, different technologies for the same modality, variability in sequencing depth, limited sensitivity, and noise. It does so while explicitly modeling the statistical properties of each modality, namely discrete signal for scRNA-seq and a largely binary signal for scATAC-seq.

Importantly, a recent method (Cobolt; available as a preprint [9]) presented an approach similar to that of MultiVI, with promising results. However, its functionality is limited to the first level of analysis (creation of a joint latent space). In the following, we utilize several published datasets to benchmark MultiVI as a more comprehensive solution for integrating and interpreting information across different modalities, studies, and technologies. In addition to showcasing its ability to derive accurate low dimensional representations, we also demonstrate several key properties of MultiVI as a way of imputing the high dimensional data. First, we demonstrate that MultiVI provides useful estimates of the uncertainty in the imputed values (i.e., predicted chromatin accessibility for scRNA-seq only cells and predicted gene expression for scATAC-seq only cells), whereby less accurate predictions are also less confident. Second, we demonstrate that these estimations of uncertainty give rise to accurate estimation of differential gene expression or chromatin accessibility in cells for which the respective modality was not available. Finally, we show that even if a population of cells has information from only one modality, accurate imputation may still be achieved when multi-modal information is available for related populations (thus effectively performing out-of-sample prediction). MultiVI is available in scvi-tools as a continuously supported, open source software, along with detailed documentation and a usage tutorial https://docs.scvi-tools.org/.

## 2 Results

### 2.1 The MultiVI Model

MultiVI leverages our previously presented variational autoencoding (VAE [10]) models for gene expression (scVI [11]) and chromatin accessibility (PeakVI [12]). Given multi-modal data from a single cell (*X*) from sample (or batch) *S*, we divide the observations into gene expression (*X*_*R*_) and chromatin accessibility (*X*_*A*_). Two deep neural networks termed ‘encoders’ learn modality-specific, batch-corrected multivariate normal distributions that represent the latent state of the cell based on the observed data, *q* (*z*_*R*_|*X*_*R*_, *S*) and *q* (*z*_*A*_|*X*_*A*_, *S*), from the expression and accessibility observations, respectively. To achieve a latent space that reflects both modalities, we penalize the model so that the distance between the two latent representations is minimized and then estimate the integrative cell state *q* (*z*|*X*_*R*_, *X*_*A*_, *S*) as the average of both representations. For “unpaired” cells, i.e., cells for which only one modality is available, the cell state is drawn directly from the representation for which data is available (i.e., *z*_*R*_ or *z*_*A*_).

In the second part of the model, observations are probabilistically generated from the latent representation using two modality-specific deep neural networks termed ‘decoders’. Similar to our previous generative models for gene expression (scVI) and accessibility (PeakVI), the model assumes that the RNA expression data is drawn from a negative binomial distribution, and the accessibility data fits a Bernoulli distribution. The likelihood of the model is computed from both modalities for paired (multi-modal) cells, and only from the respective modality of unpaired cells. Finally, during training, we include an adversarial component which penalizes the model if cells from different modalities are overly separable in latent space.

This two-part architecture enables MultiVI to achieve several goals: first, it leverages the paired data to learn a low-dimensional representation of cell state, which reflects both data types. Second, it allows cells for which only one modality is available to be represented at the same (joint) latent space. Finally, the ‘decoding’ part of the model provides a way to derive normalized, batch-corrected gene expression and accessibility values for both the multi-modal cells (i.e., normalizing the observed data) and for unpaired cells (i.e., imputing unobserved data; see Supplemental Figure 1 and methods).

### 2.2 MultiVI integrates paired and unpaired samples

To study how well MultiVI integrates paired and single-modality data into a common low-dimensional representation, we inspected the outcome of artificially unpairing a jointly profiled dataset. Using a multi-modal peripheral blood mononuclear cells (PBMC) dataset from 10X genomics, a randomly selected set of cells (at varying rates, from 1% to 99%), are made unpaired such that each cell appears twice: once with only gene expression data, and once with only chromatin accessibility data. This action resulted in a heterogeneous dataset containing three sets of cells: one set has both modalities available, a second set has only RNA-seq information, and the third set of cells has only ATAC-seq information present.

We compared MultiVI to Cobolt ([9]), a model similar to MultIVI that uses products of experts to create a common latent space. To explore the performance of additional analysis strategies and since to the best of our knowledge there are no published methods for integrating multi-modal data with single-modality data, we also added several adaptations of Seurat [13]. Specifically, we attempted to use the Seurat V4 code base with three different approaches: (1) *gene activity*: we converted the ATAC-seq data of the accessibility-only cells to gene activity scores (using the *signac* procedure), and then integrated all the cells using the gene-level data (i.e., gene scores when RNA-seq is not available, or gene expression when RNA-seq is available); (2) *imputed*: we followed the steps in (1) and then used Seurat to impute the RNA expression values for the accessibility-only cells. This is done by averaging over nearby cells in the integrated space for which RNA-seq is available (methods). The data from the accessibility-only cells was then re-integrated with the remaining cells using the imputed RNA expression values instead of the gene scores; (3) *WNN*: using weighted nearest neighbor graphs, which leverages information from both modalities to create a joint representational space, then project single-modality data onto this space (methods).

We ran all methods on the artificially unpaired datasets and compared their latent representations (with the exception of application of the WNN-based approach on the 99% unpaired dataset, which failed to produce results due to the low number of paired cells; Figure 1A-C, Supplemental Figure 2). We first quantified the mixing abilities of the different approaches, by calculating the local inverse Simpson’s index (LISI) score described by [14]. Briefly, for each unpaired cell the fraction of neighbors among the K nearest single-modality neighbors that are of the same modality (expression or accessibility), for varying values of K, normalized by the overall fraction of that modality. This results in an enrichment score, with 1 being perfect mixing (Figure 1D). We found that algorithms based on generative modeling (Cobolt, MultiVI) outperform the alternative approaches of gene scoring and WNN in most rates of unpaired cells. Conversely, the Seurat-based imputation approach (unlike the other two Seurat-based approaches) maintains high mixing performance across all levels of unpaired cells. This result is expected, though, since each accessibility-only cell is represented by an average of cells for which RNA-seq data is available and that have similar gene expression profiles (i.e., a local neighborhood in a transcriptome-based space). It does not, however, indicate whether these representations are accurate.

**Figure 1:**
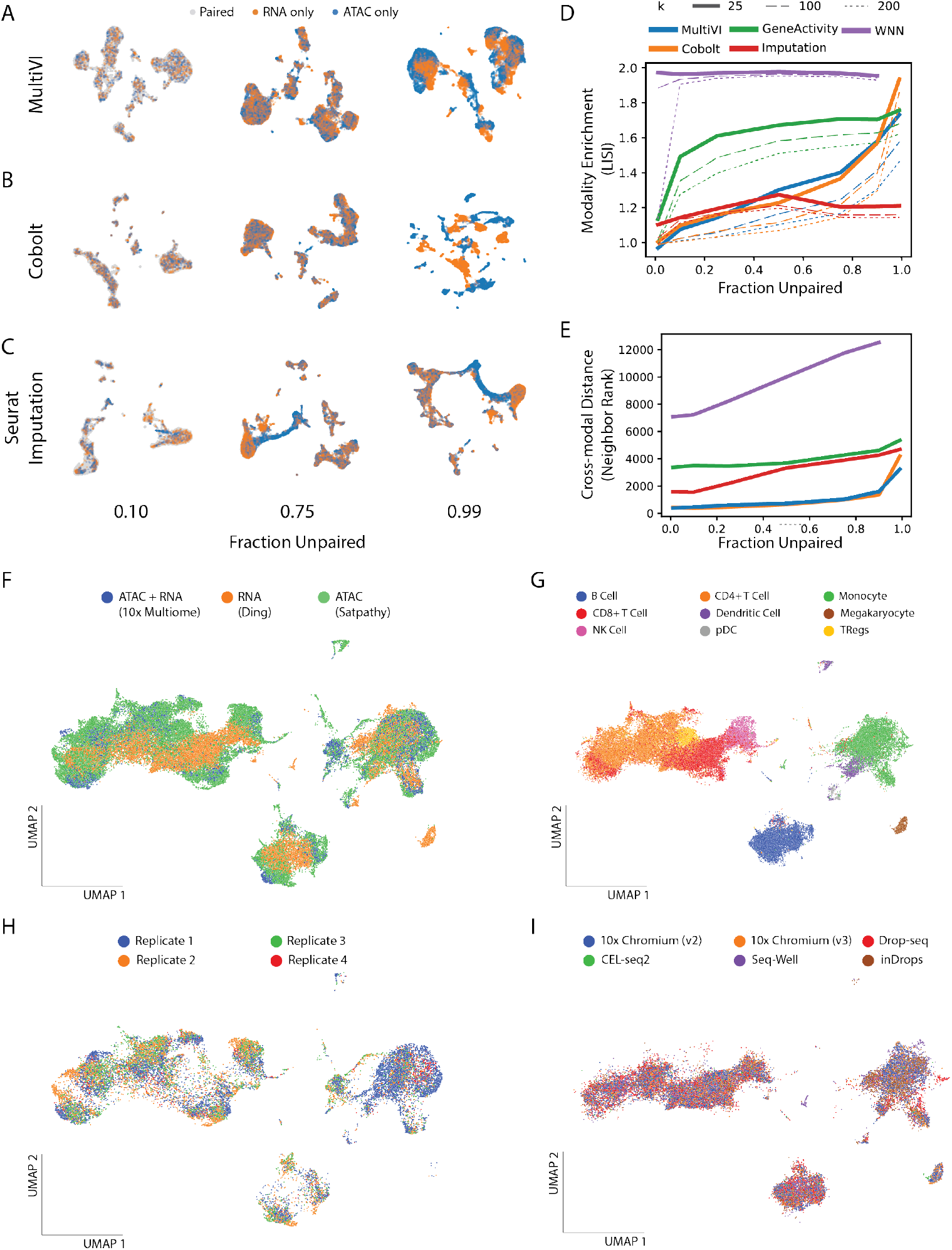
MultiVI accurately integrates gene expression and chromatin accessibility data. A-C) UMAP representations of the latent spaces learned by MultiVI (A), Cobolt (B), Seurat using the RNA-imputation based integration (C), for various rates of unpaired data, colored by cell modality. D) Modality Enrichment (LISI score), computed as the fraction of neighbors of the K-nearest neighbors that are from the same modality, normalized by the overall fraction of the cells from that modality. E) The mean distance between the two representations of artificially unpaired cells, measured as the number of cells between them. F-I) UMAP representation computed from the latent space of MultiVI in which cells are color labeled by: (F) their modality; (G) cell type label; (H) scATAC-seq PBMC cells labelled by the replicate from which they were collected; (I) scRNA-seq cells labelled by their experimental technology.

To explore this, we next examined the accuracy of the inferred latent space. To measure how well each method captures the true biological state of a cell, we took advantage of the ground truth information contained in our artificially unpaired datasets. For the unpaired cases, we have two distinct representations of the same cell: one based solely on the expression profile and the other solely on the chromatin landscape. Ideally, the two representations would be situated closely in the latent representation, as both capture the same biological state. To measure this, we looked at the distances between the two representations of the unpaired cells in the latent space created by each method. To account for the varying scales of different latent spaces, we used the rank distance (the minimal *K* for which the two representations are within each other’s K nearest neighbors, averaged across all cells; methods, Figure 1E). In this experiment, we found that MultiVI and Cobolt maintain the multi-modal mixing accuracy substantially better than the three alternatives, and that all methods have a deteriorating performance as the level of unpaired cells increases.

Taken together, these results show that the deep generative modeling approach, as taken by MultiVI, efficiently integrates unpaired scRNA and scATAC data while capturing the true biological state of each cell. They also demonstrate that the alternative approaches we implemented with Seurat either mix the modalities less effectively, or mix them well but less accurately.

### 2.3 Integration of Independent Datasets

Our previous analyses rely on artificially unpaired data, where our model benefits from all data fundamentally being generated in a single batch and by a single technology. While allowing for more accurate benchmarks, this does not reflect real-world situations in which it is desired to integrate datasets that were generated at different batches or even different studies, while possibly using different modalities and technologies. We therefore sought to demonstrate MultiVI on a set of real-world data. We collected three distinct datasets of PBMCs: 1) Multi-modal data from the 10X dataset we used previously; 2) ATAC-seq from a subset of Hematopoeisis data generated by Satpathy et al[15], containing multiple batches of PBMCs as well as cell-type specific (FACS-sorted) samples; 3) PBMC data generated by several different technologies for single cell RNA sequencing, taken from a benchmarking study by Ding et al [16]. The datasets were processed to create a set of shared features (genes or genomic regions, when measurements are available), and annotations were collected from both Satpathy et al and Ding etl al datasets and combined to a shared set of cell type labels (methods). The resulting dataset has 47148 (53%) ATAC-only cells from Satpathy et al, 30495 (34%) RNA-only cells from Ding et al, and 12012 (13%) jointly profiled cells from 10X.

To gauge the extent of batch effects in this data, we first ran MultiVI without accounting for the study of origin of each sample or to its specific technology (which varies between the RNA-seq samples from Ding et al). With this application, we found substantial batch effects, both between different samples in the chromatin accessibility data and between technologies in the gene expression data (Supplemental Figure 4). We then reanalyzed the data, this time configuring MultiVI to correct batch effects and technology-specific effects within each dataset (methods). The resulting, corrected, joint latent space mixes the three datasets well (Figure 1F), while accurately matching labeled populations from both datasets (Figure 1G). MultiVI achieves this while also correcting batch effects within the Satpathy data and technology-specific effects within the Ding data (Figures 1H-I). To better examine the correctness of the integration, we examined the set of labelled cells from the two single-modality datasets (FACS-based labels from Satpathy and manually annotated cells from Ding). For each cell, we examined its 100 nearest neighbors that came from the other modality, and summarized the distribution of labels of those neighbors. We find a clear agreement between the labels of each cell and the labels of it’s neighbors, with some mixing among related cell types (Supplemental Figure 5). Using a similar set of experiments, we also observed that the low dimensional representation inferred by Cobolt achieves similar levels of mixing and accuracy (data not shown). This analysis therefore demonstrates that MultiVI, and more generally, the deep generative modeling approach are capable of deriving biologically meaningful low dimensional representations that effectively integrate data not only data from different modalities, but also from different labs and technologies.

### 2.4 Probabilistic Data Imputation with Estimated Uncertainty

The generative nature of MultiVI enables several functionalities for analyzing the data in the full high dimensional space, i.e imputation of missing observations and modalities, estimation of uncertainty, and differential analysis. These functionalities are currently unique to MultiVI, and are not implemented by Cobolt or other generative models. To demonstrate MultiVI’s imputation abilities, we resorted to the 10X PBMC dataset where 75% of the cells were artificially unpaired (as in Figure 1). We used MultiVI to infer the values of the missing modality for the unpaired cells and found that for both modalities, the imputation had high correspondence to the observed values (Figures 2A-C). Specifically, we observe Spearman correlation of 0.57 between the imputed expression values and the observed data (taking the raw values, scaled by library size), and an area under the precision-recall curve (PRAUC) of 0.41 for the accessibility data (taking the raw, binary signal). Since the raw data can be largely affected by low sensitivity, we also calculated the correlation between the imputed values and a smoothed version of the data (obtained with a method different of MultiVI; methods), where the signal is average over similar cells (separately for ATAC-seq and RNA-seq), thus mitigating this issue. As expected, we see a higher level of correspondence between the imputed values and this corrected version of the raw data (Spearman correlations 0.8 and 0.86 for accessibility and expression respectively; Supplemental Figure 6A-B).

**Figure 2:**
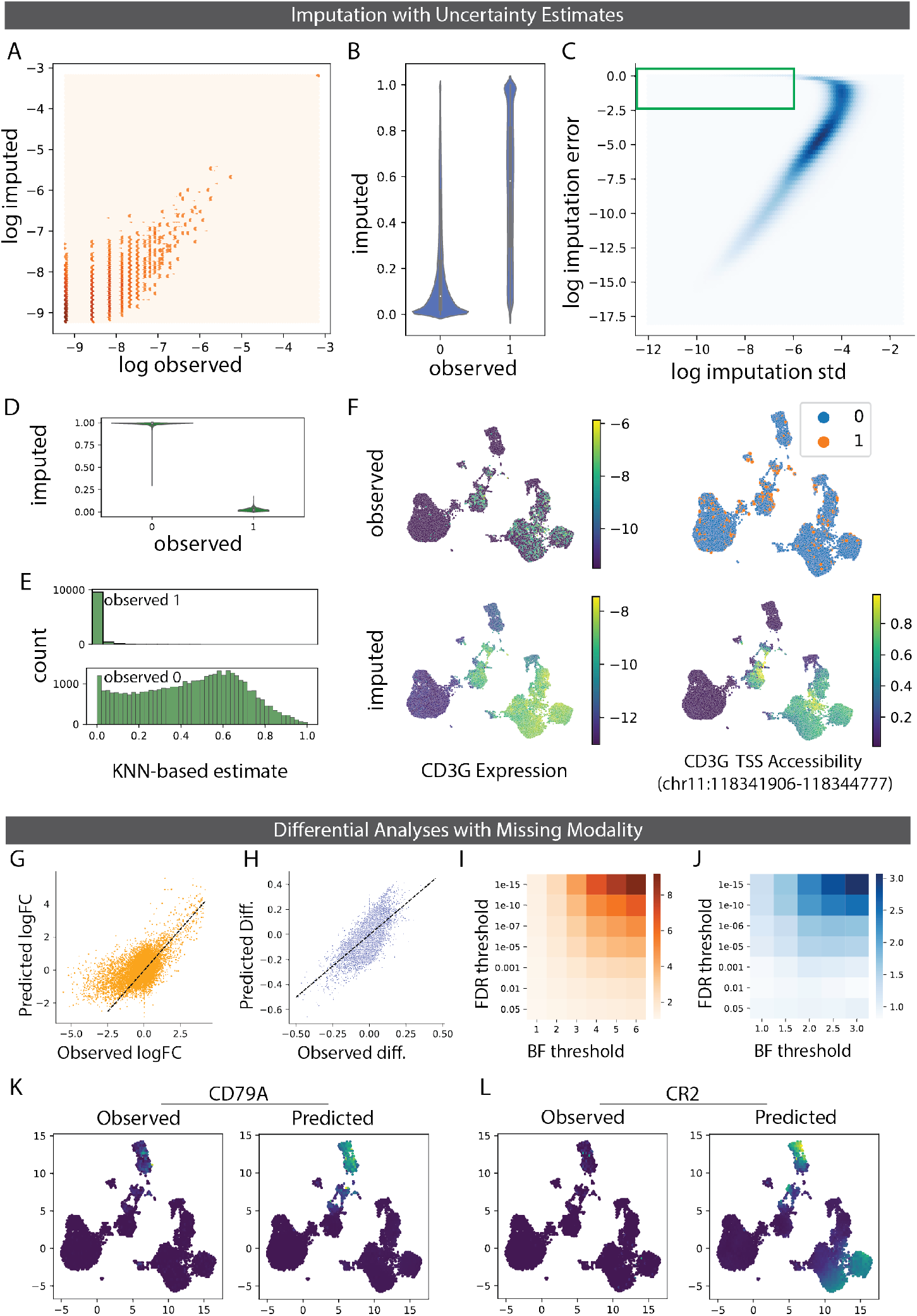
MultiVI imputation and uncertainty estimation. A) normalized observed RNA counts by MultiVI-imputed RNA estimates; all values, including color intensity, are presented in log scale (*log* (*x* + 1*e* − 4) for stability. B) MultiVI-imputed accessibility estimates by the observed values. C) the imputation error (imputed − observed)^2^ as a function of the standard deviation of the imputed accessibility estimates. Green box marks high-confidence-high-error values examined in following panels. D) MultiVI-imputed accessibility estimates by the observed values for high-confidence-high-error cases. E) smooth accessibility estimates for values observed as 1 (top) and 0 (bottom). estimates computed by averaging the accessibility profiles of the 50 nearest neighbors, in a 50-dimensional space computed using Latent Semantic Indexing. F) observed and imputed values for CD3G expression and CD3G TSS accessibility. Expression values are normalized per cell and displayed in log scale. G-H) Differential effect sizes between B-cells and the remained of the data, comparing the effects computed from the held-out expression data with those predicted by MultiVI, for differential expression (G) and differential accessibility (H). I-J) Enrichment of the overlap between statistically significant results for various significance thresholds for expression (I) and accessibility (J). K-L) expression values for B-cell marker CD79A (K) and B- and T-cell marker CR2 (L), observed in the held-out data (left) and predicted by MultiVI (right), displayed using latent space coordinated computed using all the available data.

Next, we focus our analysis on uncertainty estimation for the imputed accessibility values. We measured the uncertainty of the model for each imputed accessibility value by sampling from MultiVI’s generative model (methods) and found a strong relationship between the estimated uncertainty and the error of each data point ((imputed −observed)^2^), indicating that the model is indeed less certain of predictions that are farther from the unobserved “ground truth” values (Figure 2C). Equivalent analysis for expression imputations is hindered by the high correlation between the average expression and both the measured error and the uncertainty of the imputed results.

Interestingly, we identified a small subset of values (roughly 0.5% of observations) for which we have high confidence imputations that are associated with high error, when comparing to the unobserved raw accessibility data (Figure 2C, green square). In the case of chromatin accessibility, these high-confidence-high-error imputations correspond to cases where the model confidently predicts the opposite of the actually observed value (Figure 2D). To investigate the source of these errors, we inspected the same cases when comparing the imputed values to the smoothed accessibility estimates (methods). We found that many of these regions were detected as inaccessible in the raw data, but predicted to be accessible by MultiVI, and vice-versa. Interestingly, the smoothed data agrees with the MultiVI predictions - namely, observations that were predicted as accessible tend to be open in highly-similar cells, and observations that were predicted as inaccessible tend to be closed in high-similar cells (Figure 2E). This indicates that these high-confidence-high-error values may correspond to false-negatives and false-positives in the raw data.

As a specific example for imputation, we highlight the T-cell marker gene CD3G. While the observed expression and the observed accessibility of the region containing the transcription start site (TSS) of the gene show high noise and sparsity, the imputed values are highly consistent and clearly mark the T-cell compartment of the latent space (Figure 2F). Overall, these results show that MultiVI is capable of imputing missing observations, and quantifying the uncertainty for each value, allowing the user to then determine which imputed values are reliable for downstream analyses and which are not.

### 2.5 Cross-modal Differential Analyses

Our previous results demonstrate that MultiVI can be used to accurately impute missing observations of single cells, in situations where the multi-modal and the single-modality data both contain the same cellular subsets. The imputation task becomes more challenging when analyzing a population in which one of the modalities was not observed at all. However, the ability to impute values in this scenario will enable leveraging multi-modal data to analyze a wider variety of single-modality datasets, even if a fully matching multi-modal data is not available.

To explore this, we used the same 10x PBMC multiome dataset, with 75% of cells artificially unpaired, and clustered the latent space to identify distinct cellular populations (Supplemental Figure 7A). We chose the B cell cluster, which we annotated as such using established markers (e.g CD19, CD79A). Next, we corrupted the data further by removing all expression information (paired or unpaired) from the B-cell population, thus creating a distinct population for which only accessibility data is available to the model. In a second experiment, we removed all accessibility data from the same compartment to create a dataset for which only expression was observed for B-cells (Supplemental Figure 7B-C). We trained MultiVI separately on each of the two corrupted datasets, and used the model to perform differential analyses, comparing the B-cell population and the remainder of the cells. Specifically, we conducted differential expression analysis with the model trained without B-cell expression data (corrupted dataset 1), and differential accessibility with the model trained without B-cell accessibility data (corrupted dataset 2). Estimates of significance were done with a Bayes factor, as in previous work [11, 12, 17](Methods). To evaluate the accuracy of this analysis, we used standard differential analyses (not using generative models) on the held-out data to create ground-truth results and compared them to our inferred results (Methods). Considering the first corrupted dataset, although no expression data was observed in the B cell population, we found high concordance between the observed and predicted log Fold-Change values (Figure 2G, Pearson’s correlation 0.57). When examining genes that are preferentially expressed in B-cells (observed logFC ¿ 1) this became more evident (Pearson’s correlation 0.74). Similarly, with the second corrupted dataset, we found high concordance between observed and predicted differences of accessibility (Figure 2H, Pearson Correlation 0.67).

Among the top most differentially expressed genes, we found known B-cell markers, including IGLC3, IGHM, CD79A, and IGHD (Supplementary Table 1). Overall, we identified 1621 significantly differential genes (FDR[17] ¡ 0.05), of which 75% were also identified with the held-out data at a 5% false discovery rate (FDR), thus representing a modest but significant enrichment (odds-ratio 1.22, Hypergeometric test *p* < 1.9^−35^; Supplementary Table 1). Increasing the threshold of significance (on the FDR for the standard analysis, and the Bayes Factor for the MultiVI results) increased the overlap between the sets of results, indicating that the results are more consistent for more highly significant genes (Figure 2I). Similarly, we identified 922 differentially accessible regions (FDR[17] ¡ 0.05), of which 86% were also identified with the held-out data at 5% FDR (odds-ratio 1.57, *p* < 1.7^−95^). As in the expression analysis, the overlap between the inferred and observed differential accessibility analyzes increased with the significance thresholds (Figure 2J).

Finally, we predicted the expression of genes identified as preferentially expressed in B-cells by the model trained without B-cell expression data. CD79A, which encodes for part of the B-cell receptor complex, and one of the top genes identified by MultiVI, was indeed found to have highly localized predicted expression in the B-cell compartment (Figure 2K, displayed using original UMAP coordinates as in Figure 2F). Another differentially expressed gene, CR2, a membrane protein found on both B- and T-cells, was predicted specifically on the corresponding compartments (Figure 2L).

Taken together, these results demonstrate that MultiVI can be used to impute missing modalities even for populations that were only identified in a single-modality dataset. This unlocks the ability for leveraging multi-modal data to reanalyze existing single-modality datasets and impute the missing modality: chromatin landscape for existing scRNA experiments, and gene expression for existing scATAC experiments, as well as perform differential analyses using these imputed values.

## 3 Discussion

MultiVI is a deep generative model for the integrated analysis of single cell gene expression and chromatin accessibility data. MultiVI uses jointly profiled data to learn a multi-modal model of the data and to relate measurements of individual modalities on the same population of cells. The model accounts for various technical sources of noise and can correct additional sources of unwanted variation (e.g batch effects). MultiVI learns a rich latent representation of the data coalescing information present in each individual data type, which can be used for further single-cell sequencing analysis.

Recent algorithms for the analysis of multi-modal data were developed to process paired datasets, in which both modalities have been profiled at the same cell [8, 7]. These algorithms handle multi-modal data, but lack the ability to integrate single modality datasets into the same analysis. While this task is possible to achieve with the Seurat code base [13], the respective methods we utilized here were not specifically designed to this end, and their performance was not tested for this task. Here, we have shown that use of deep generative modeling, either with MultiVI or the recently presented Cobolt [9] can effectively combine unpaired scRNA and scATAC data with multi-modal single-cell data, generating a robust and meaningful representation of the cells’ state that captures information about both their transcriptome and epigenome. Importantly, this joint representation is achievable even when the amount of paired data is minimal, thus opening exciting opportunities for future studies in which only a small amount of paired data can be sufficient for deriving a more nuanced interpretation of single modality data.

An additional, key capability that is unique to MultiVI is the inference of the actual values of the missing modality. We have demonstrated here that this can be used to identify preferential gene expression in sub-populations for which only chromatin accessibility data is available and distinguishing chromatin features for sub-populations for which only gene expression data is available. These results open the way for exciting future applications. First, MultiVI and similar methods have the potential to enable a reanalysis of the large compendia of available single modality datasets (representing the majority of existing data) with relatively small additional paired data, thus potentially leading to more comprehensive characterizations of cell state. Second, it can facilitate cost-effective designs for future studies, in which only a subset of samples need to be profiled with the (more costly) multi-modal protocol.

In summary, MultiVI is able to seamlessly process single and multi-modal data, integrate different chromatin and transcriptional batches, and create a rich joint representation harnessing all available information. It is implemented in the scvi-tools framework [18], making it easy to configure, train, and use.

## Declarations

## Acknowledgements

We thank Adam Gayoso for assistance on model implementation in scvi-tools. We thank Christina Usher for assistance with visualizations.

## Author contributions

TA, MIG and NY conceived of the model and designed the analyses. TA and MIG implemented the model with input from AG. TA performed the analyses. NY and MJ supervised the work. TA, MIG, MJ and NY wrote the manuscript.

## Data and Software Availability

All data used in this manuscript is publicly available via the original publications and releases. Intermediate data, trained models used in this manuscript, and the notebooks to generate the figures in this manuscript, are all posted and available on zenodo: 10.5281/zenodo.5225858.

## 4 Methods

### 4.1 The MultiVI Model

MultiVI inherits generative models describing chromatin accessibility and transcriptional observations from scVI [11] and peakVI [12]. Briefly, Let 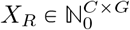 be a scRNA-seq genes-by-cell matrix with *C* cells and *G* genes, where 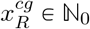 is the number of reads from cell *c* that map to gene *g*. Let 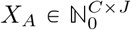 be a scATAC-seq region-by-cell matrix with *C* cells and *J* regions, where 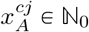 is the number of fragments from cell *c* that map to region *j*.

MultiVI models the probability of observing *x*_*cj*_ counts in a gene by using a negative binomial distribution,

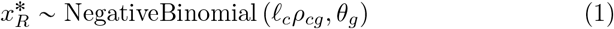

where *ℓ*_*c*_ is a scaling factor that captures cell-specific biases (e.g library size), *ρ*_*c*_*g* is a normalized gene frequency and *θ*_*g*_ models the per gene dispersion. The probability of observing a region as accessible is modeled with a Bernoulli distribution,

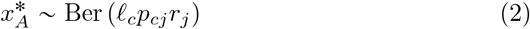

where *p*_*cj*_ captures the true biological heterogeneity; *r*_*j*_ captures region-specific biases (e.g width, sequence). In both observational models, the scaling factor the region-specific and the per gene dispersion parameters are inferred from data using deep neural networks (this is in contrast to the original implementation of SCVI in which library size was modelled using a lognormal distribution).

Next, for each cell, normalized gene frequencies *ρ*_*cg*_ and biological heterogeneity *p*_*cj*_ are estimated using a latent representation as in VAE[10]. Briefly, each modality is assign their own latent representation, a isotropic multivariate normal distribution 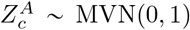 and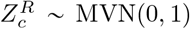. Then, with the purpose of bringing both representations together, they are combined by taking their average 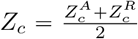. This merged representation is then used to decode both model parameters, *p*_*c*._ = *f* (*Z*_*A*_) and *ρ*_*c*,_ = *g*(*Z*_*A*_).

### 4.2 MultiVI Inference Model

We use variational inference [19] to compute posterior estimates of model parameters using the following variational approximation:

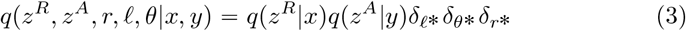

where delta distribution *δ* highlight the fact that parameters are inferred from the data as point estimates. The cell-specific factor *ℓ*_*c*_ is computed from the input data for cell *c* via a deep neural network 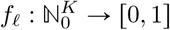. The region-specific factor *r*_*j*_, since it is optimized across samples, is stored as a *K*-dimensional tensor, used and optimized directly. In the case of each latent representation, two encoders are computed as 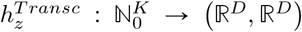 and 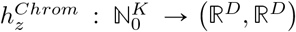 where each of them computes the distributional parameters of a D-dimensional multivariate normal random variable: *Z* ⋃ *MVN*((*h*_*z*_ (*x*_*c*_)_1_, *h*_*z*_ (*x*_*c*_) _2_).

Using the variational approximation, the evidence lower bound (ELBO) is computed and optimized with respect to the variational and model parameters using stochastic gradients. To enforce the similarity between chromatin and transcription latent representations, we add to the ELBO a term that penalizes the distance between representations using a symmetric KL divergence between distributions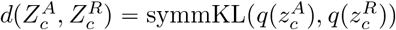.

### 4.3 Modeling differences between MultiVI and Cobolt

While conceptually similar, MultiVI and Cobolt have several key differences in design and implementation choices. MultiVI offers additional functionalities due to its generative model, i.e denoising, imputation, uncertainty estimation, and differential analyses - are discussed in detail in this manuscript. In addition to those, we detail several other differences between the methods: (a) MultiVI uses a distributional average and penalization to mix the latent representations, compared with the classical product of experts calculation used by Cobolt. (b) the distributional assumptions made by the models are different: MultiVI uses tailored noise models for each modality (negative binomial for expression, Bernoulli for accessibility), and uses a deep neural network for the generative component of the model as well as the inference component. In contrast, Cobolt uses a multinomial likelihood for both modalities and uses a linear transformation as a generative model. (c) MultiVI explicitly avoids over-fitting the data, in both the architecture (e.g droplout layers) and training procedure (holding out data to use for early-stop if the model overfits), whereas Cobolt does not contain such guardrails.

## 5 Benchmarking and Evaluation

### 5.1 Dataset Preprocessing

The 10x multiomic unsorted PBMC dataset was downloaded from the company website. For artificial unpairing analyses, the processed peak-by-cell matrix was downloaded and filtered to remove features that are detected in fewer than 1% of the cells. For the mixed-source PBMC dataset, the fragment file was downloaded and reprocessed using CellRanger-ARC (v2.0.0) with the Satpathy hg38 peaks. The Satpathy dataset was downloaded from GEO (Accession GSE129785); specifically the processed peak-by-cell matrix and meta-data files: *scATAC*−*Hematopoiesis All*.*cell*−*barcodes*.*txt*.*gz, scATAC*−*Hematopoiesis*−*All*.*mtx*.*gz, scATAC*−*Hematopoiesis*−*All*.*peaks*.*txt*.*gz*. We then filtered the data to only include peaks that were detected in at least 0.1% of the data, and lifted those peaks over from the hg19 to the hg38 genome reference using the UCSC liftover utility [20]. The Ding dataset was downloaded from GEO (Accession GSE132044); specifically the pbmc data: *pbmc*_*h*_*g*38_*c*_*ount*_*m*_*atrix*.*mtx*.*gz, pbmc*_*h*_*g*38_*c*_*ell*.*tsv*.*gz, pbmc*_*h*_*g38*_*g*_*ene*.*tsv*.*gz*.Matching cell type annotation was downloaded from SCP (Accession SCP424). After preprocessing, the reanalyzed 10x dataset was combined with both single-modality datasets, and the features were filtered to remove features (either genes or peaks) that were detected in fewer than 1% of the cells.

### 5.2 RNA-based Seurat integration

This integration modality, disregards multiomic information and only RNA information is considered from multiome cells. Briefly, RNA information is first integrated and then, chromatin accessibility is integrated using gene activity scores (*RNA-based* method) or RNA imputed values (*RNA-based Imputed* method).

In more detail, cells were separated into three different datasets, multiomic cells (using only expression data), rna-only cells and atac-only cells. Seurat objects were created for multiome and rna-only data, and were then normalized, scaled, and the first 50 principal components are calculated. For atac-only cells, a Seurat object was created, gene activity scores were calculated, scaled, and principal components were computed. To integrate the three datasets, integration anchors (using *FindIntegrationAnchors*) were calculated and the data was then integrated (using *IntegrateData*). The *RNA-based* method uses gene activity scores as representative values from the atac-only cells. The *RNA-based Imputed* method includes an additional step in which RNA imputed values are calculated from gene activity scores by running *FindTransferAnchors* and *TransferData*.In this integration method, RNA imputed values are used as representative values from atac-only cells. Finally, integrated data was then scaled and principal components were calculated to generate the final latent space. Across these integration methods, we followed the standard recommended procedure for analyzing data with Seurat given in their tutorials [21].

### 5.3 WNN-based Seurat Integration

This approach aims to leverage information from both modalities (chromatin accessibility and expression values), using the newly described weighted nearest neighbors approach from Seurat V4 [13]. We first created a weighted nearest neighbor graph using multiomic information and then project chromatin and transcriptional information onto this.

We begin by separating cells in unpaired datasets into three different datasets, multiomic cells (with both expression and chromatin data), rna-only, and atac-only. First, multiome latent representation is found by calculating SC transform and principal components on the expression data and latent semantic analysis (TF-IDF decomposition followed by SVD) on the chromatin data. Next, multimodal neighbors and the first 50 supervised PCA are calculated. To merge rna only and atac only data to multiome representation, transfer anchors (FindTransferAnchors) are computed on rna only data and gene activity scores on atac only and each datasets is integrated using IntegrateEmbeddings function. Finally, datasets and dimensionality reductions are merged and umap is visualized using the merged information.

### 5.4 Neighbor Rank Distance Calculation

For artificially unpaired cells, each cell has two unpaired representations in the latent space. Given cell *c* with representations *c*_*a*_ and *c*_*b*_, let *S* (*c*_*a*_, *K*) be the set of K nearest neighbors to *c*_*a*_. We then define *δ* (*c*_*a*_, *c*_*b*_) as the minimal *K* for which cell *c*_*b*_ is among the K nearest neighbors of cell *c*_*a*_: min {*k*: *c*_*b*_ ∈ *S* (*c*_*a*_, *k*)}.

### 5.5 LISI score Calculation

Enrichment scores were computed as they were in our previous work[12], and similarly to the LISI scores described in the Harmony paper[14]. Briefly, given a latent representation *R*, an integer *k*, and the modality labels (expression, or accessibility) *L*, we compute *G*_*R,k*_ the K-nearest neighbor graph from *R* with *k* neighbors. Using *G*_*R,k*_, we compute for each cell the proportion of neighbors that share the same modality: 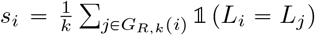. The enrichment score is the average score across all cells, 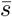, normalized by the expected score for a random sample from the distribution of labels: 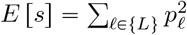, with *p*. ℓ being the proportion of each modality.

### 5.6 Estimating Imputation Uncertainty

We estimate the uncertainty of the model for each imputed value by sampling from the latent space (n=15) and computing the standard deviation of the imputed values for each observation. More consistent predictions correspond to less uncertainty.

### 5.7 KNN-based estimate of accessibility

To estimate accessibility without using MultiVI, we computed a lower-dimensional representation of the data using Latent Semantic Analysis (LSA, top 30 components), then for each cell we computed the average accessibility profile of the 50 nearest neighbors in the LSA space. This creates a smooth estimate of accessibility using highly-similar cells, mitigating the effect of false observations.

### 5.8 Expression Smoothing

Expression smoothing was achieved by taking the top 30 principle components of the expression data (computed with PCA), computing the K-nearest neighbors graph (for *K =* 50) and averaging the expression values of the neighbors for each cell (scaled by library size).

### 5.9 Differential Analyses with held-out data

To identify a distinct population of cells, we used the Leiden community detection algorithm[22], then examined the expression levels of known marker genes (CD79A, CD19) to identify the cluster of B-cells. We then unpaired the data within the cluster, once by removing all expression data from the B-cells and once by removing all accessibility data from the clusters. Since the data was already unpaired, this resulted in several cells with no observations at all, and those were removed from the dataset.

### 5.10 Differential Expression using held-out data

Differential expression was computed in two ways: 1) using the held-out data, values were normalized per-cell by dividing the expression levels by the total number of reads in the cell. log Fold-Change values were then computed by dividing the mean expression values in the two groups. Statistical significance was determined using Wilcoxon rank-sum test. 2) without the held-out data, using MultiVI, in a procedure described by Lopez et al[11] which samples from the latent space and uses the generative model to estimate expression profiles. Statistical significance was then determined using Bayes Factors, as well as an FDR approach described by Lopez et al[17].

### 5.11 Differential Accessibility using held-out data

Differential accessibility was computed equivalently to differential expression. 1) using held-out data, values were normalized using the TF-IDF transformation, differential accessibility was computed by subtracting the mean accessibility in the reference group from the same value in the target group. Statistical significance was determined using Wilcoxon rank-sum test. 2) Without the held-out data, using MultiVI, using the procedures described in our previous works[11, 12, 17].

## 6 Figures

**Figure S1:**
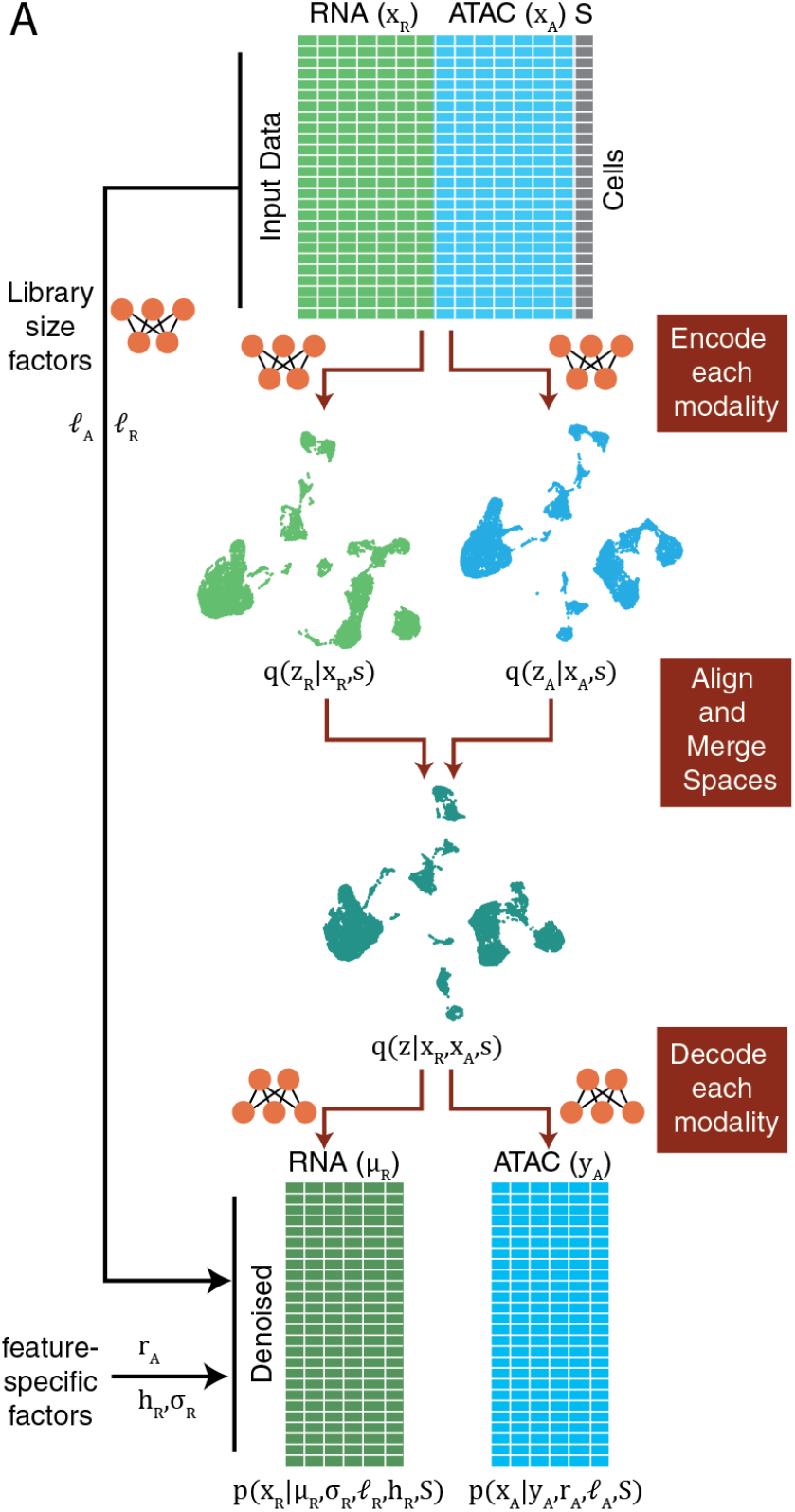
MultiVI Model Overview. Conceptual model illustration in which input data (top) consists of either chromatin accessibility (ATAC), gene expression (RNA) or both data types (Multiome). Variable *S* represents experimental covariates, such as batch or experimental condition. Each data modality is encoded into modality-independent latent representations (using neural network encoders) and then, these representations are merged into a joint latent space. The joint latent representation is used to estimate (decode) the input data together with chromatin region-specific effects (*r*_*A*_), gene-specific dispersion (*σ*_*R*_), cell-specific effects (*ℓ*_*A*_, *ℓ*_*R*_), accessibility probability estimates (*Y*_*Z*_) and mean gene expression values (*µ*_*R*_).

**Figure S2:**
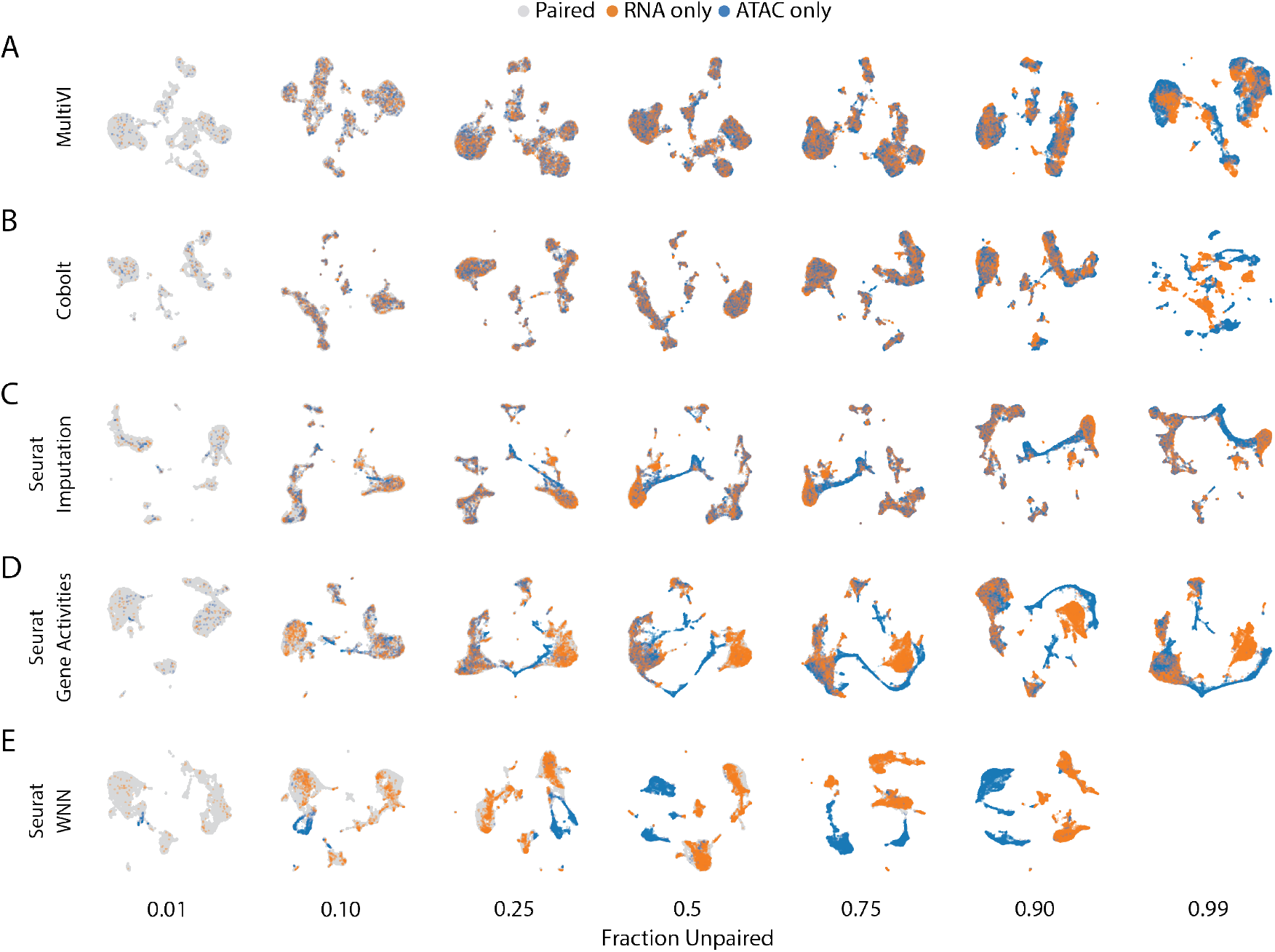
Extended Integration results depicting mixing of cells in data sets with different fraction of cells unpaired. UMAPS of latent representations for MultiVI (A), Seurat imputation method (B), Seurat Gene Activity Scores method (C), and Seurat wKNN method (D).

**Figure S3:**
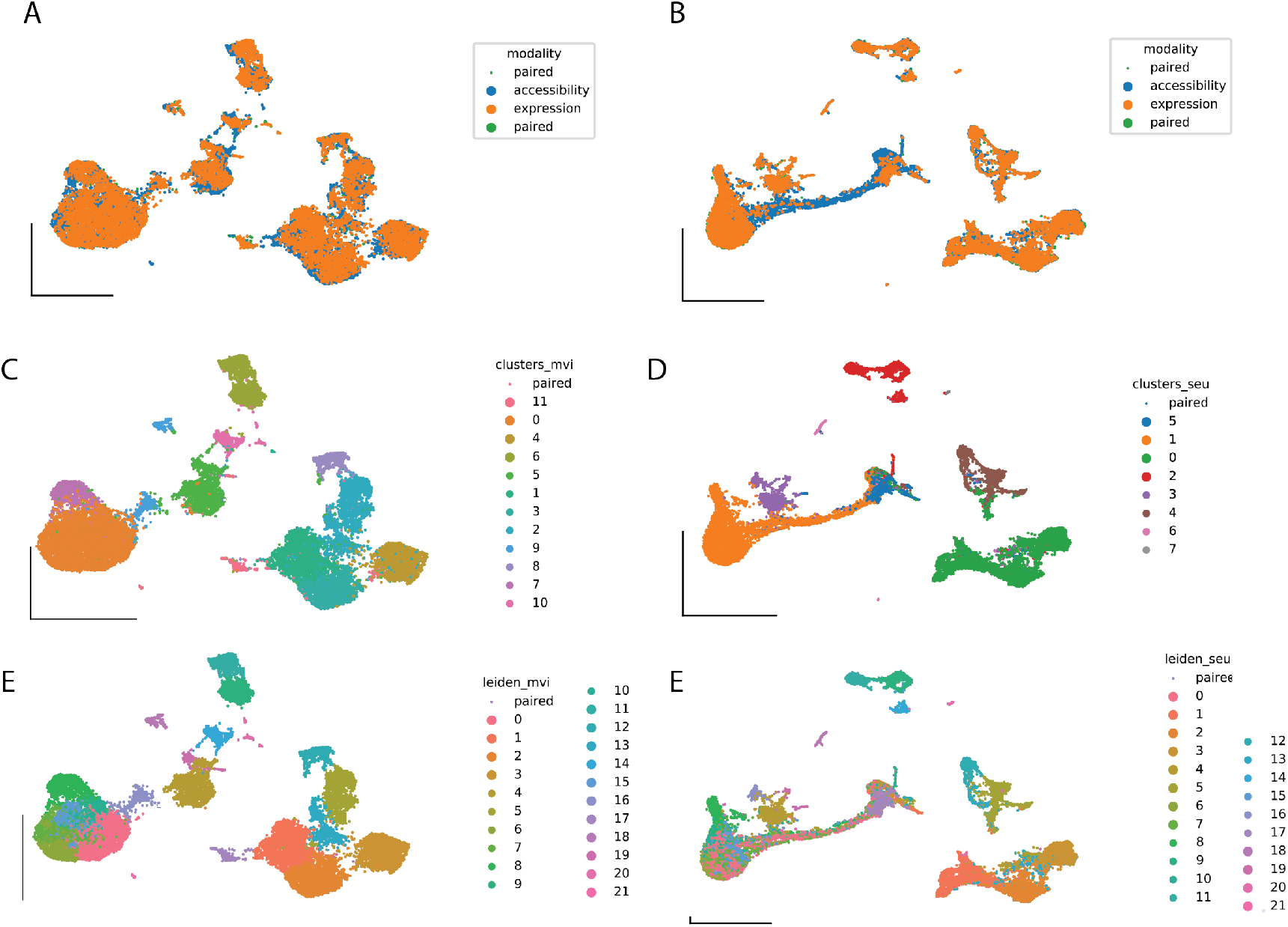
Extended Analysis of cluster consistency using dataset with 0.75 fraction of unpaired cells. UMAP representation computed from the latent space of MultiVI or Seurat Imputation in which cells are color labeled by their modality (A / B), their cluster correspondence computed at 0 fraction of unpaired cells (C / D) and their cluster correspondence computed at 0.75 fraction of unpaired cells (E / F).

**Figure S4:**
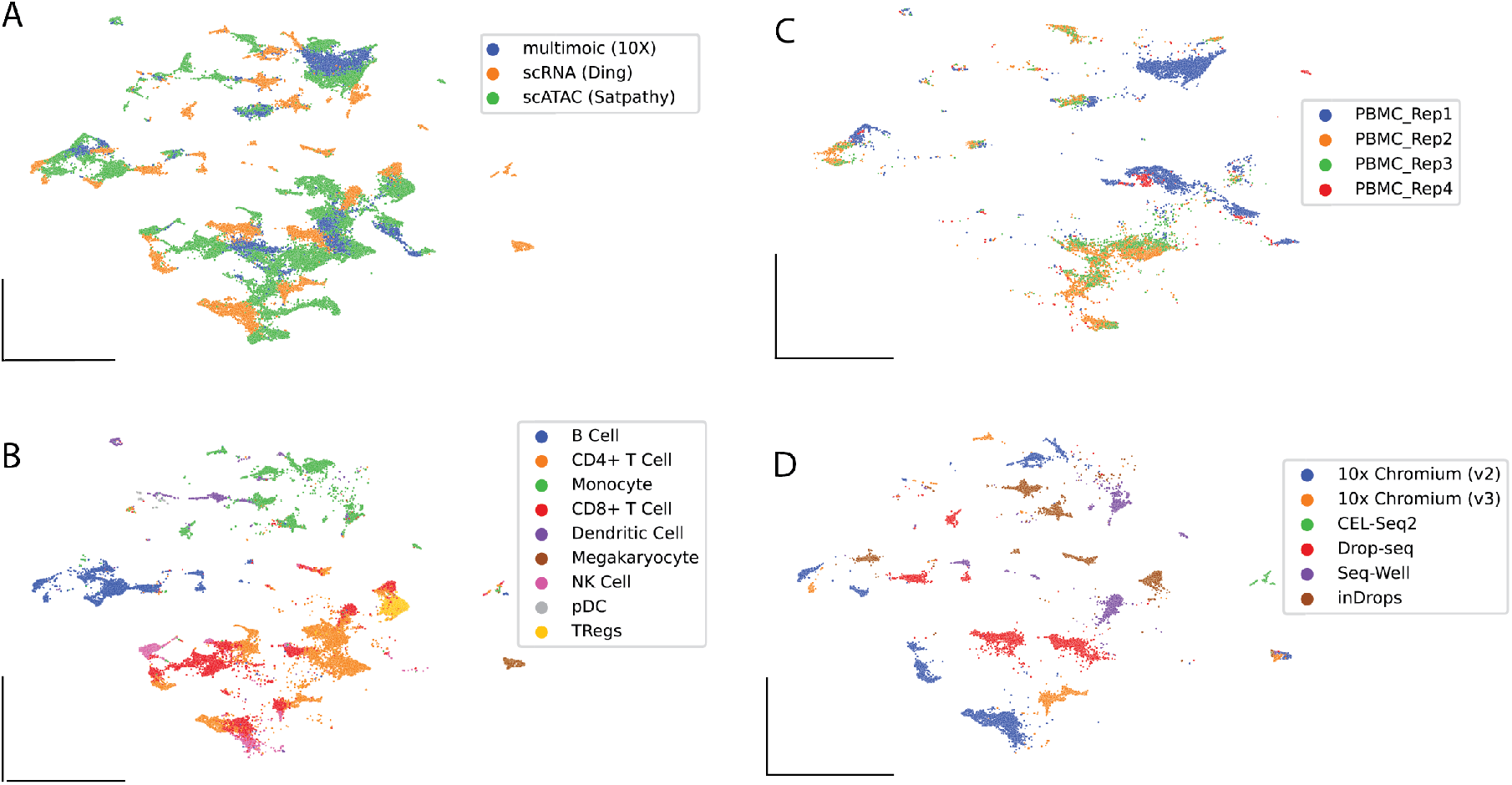
Latent representation of mixed sources data sets in which no batch correction techniques have been applied. We integrated three PBMC datasets in which only multi-modal data (10X multiome), only ATAC-seq information (Satpathy et al) and only RNA-seq information (Ding et al) is present without correcting for batch or modalities effects. A)-D) UMAP representation computed from the latent space of MultiVI in which cells are color labeled by their dataset (A), their cell type (B) or ATAC-seq cells are labelled by the replicate in which they were collected (C) or RNA-seq cells are labelled by their collection experimental technology.

**Figure S5:**
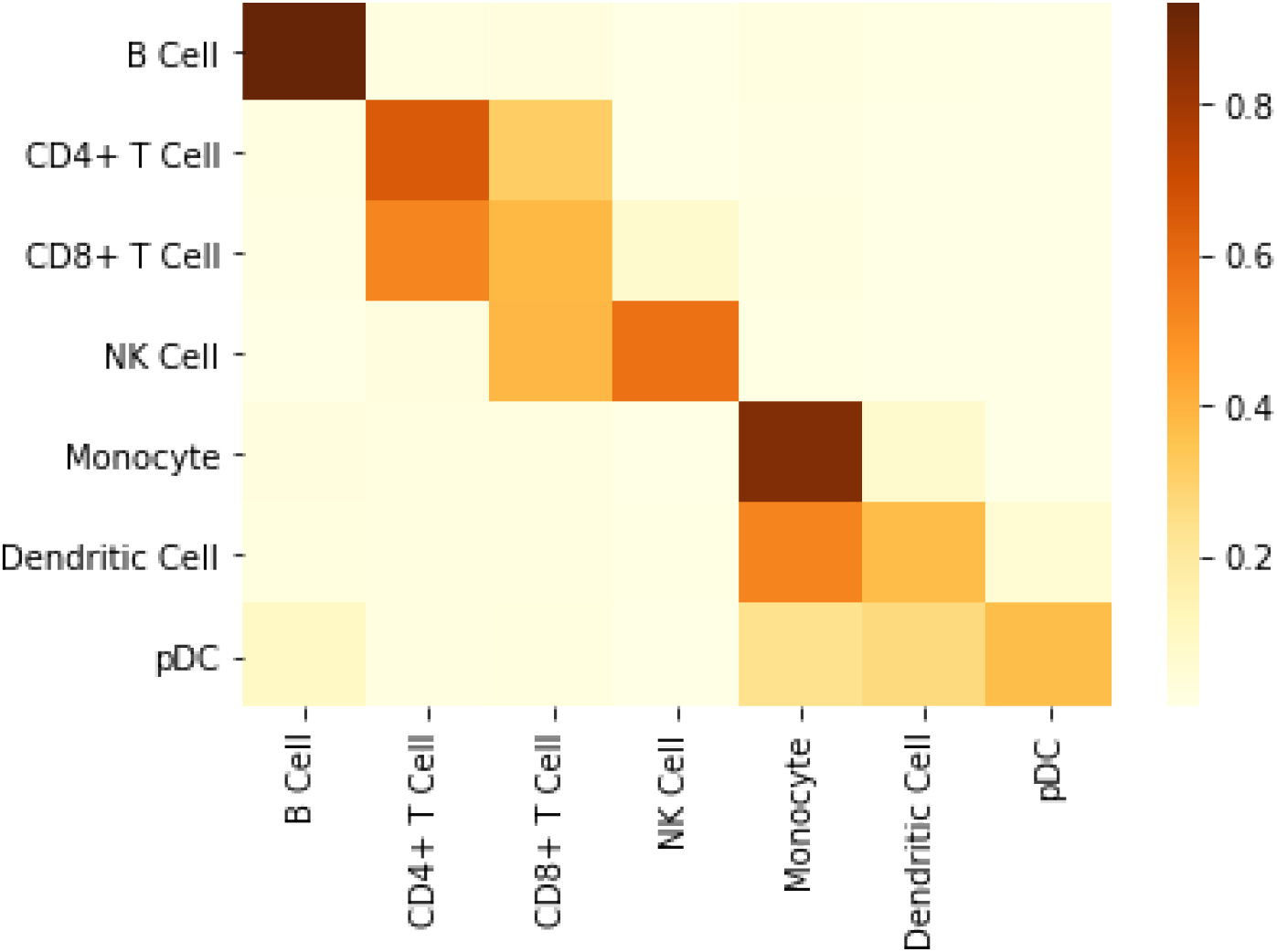
The distribution of labels of neighboring cells by the label of origin

**Figure S6:**
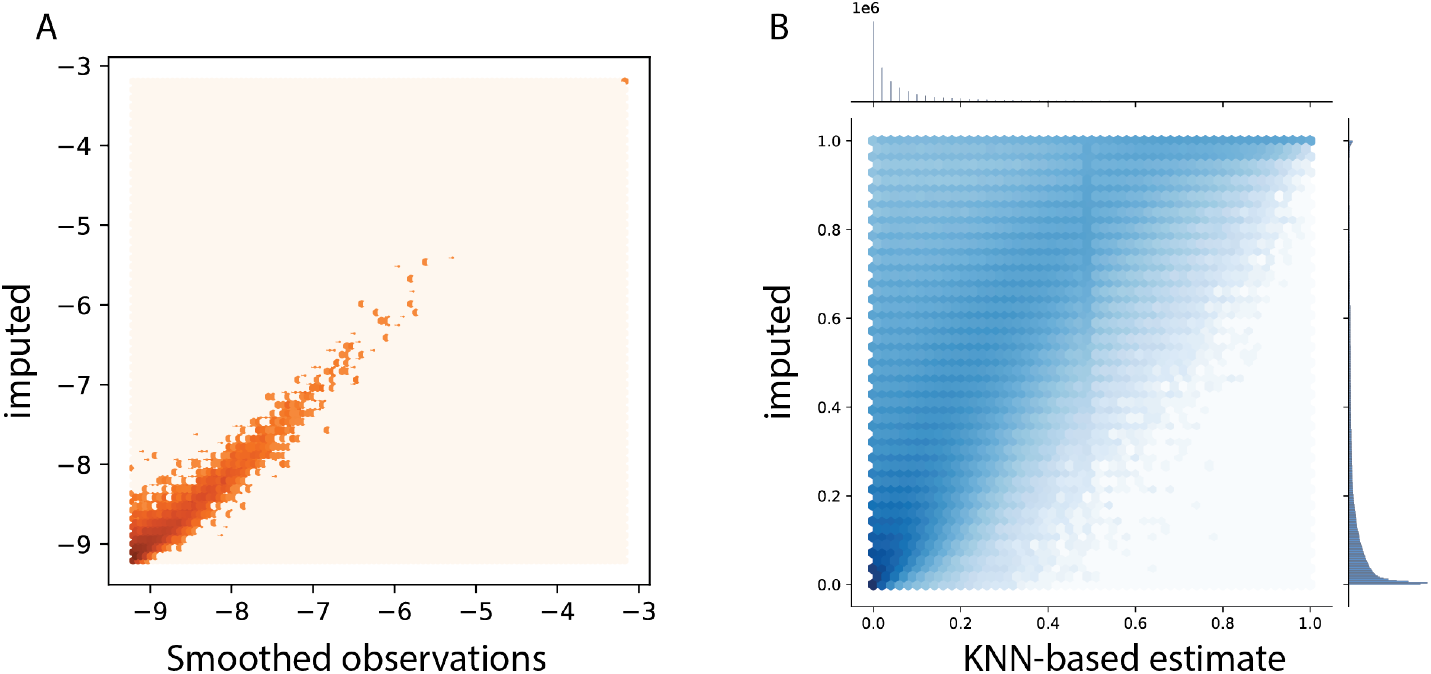
Imputed values compared with smoothed observations. Smooth averages of highly-similar cells (using 50 nearest neighbors in an independent low-dimensional space, computed separately for RNA and ATAC data) plotted against the MultiVI-imputed values for expression (A) and accessibility (B).

**Figure S7:**
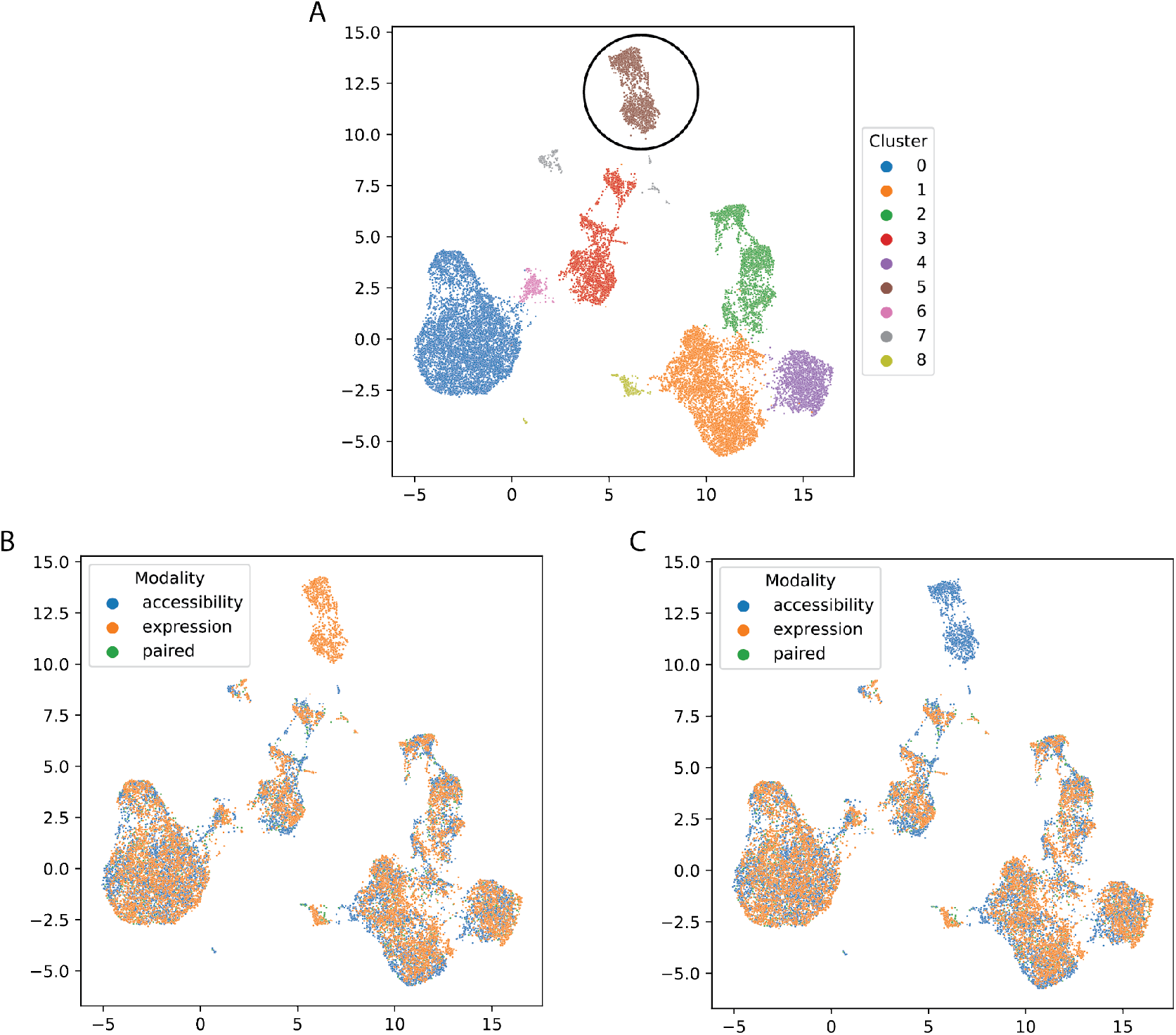
UMAP visualizations of the 10x PBMC multiome dataset, with 75% of cells artificially corrupted. A) Leiden clustering of the cells. B-C) modalities of the different cells after removal of all accessibility (B) or expression (C) data from the B-cell compartment (cluster 5).

